# Brain–heart interactions underlying differential recovery after transient systemic anoxia

**DOI:** 10.64898/2026.04.17.719210

**Authors:** Diego Candia-Rivera, Sofia Carrion-Falgarona, Mario Chavez, Fabrizio de Vico Fallani, Stéphane Charpier, Séverine Mahon

**Affiliations:** Sorbonne Université, Institut du Cerveau - Paris Brain Institute - ICM, CNRS, Inria, Inserm, AP-HP, Hôpital de la Pitié Salpêtrière, F-75013, Paris, France

## Abstract

Global cerebral anoxia is a leading cause of death and resuscitated patients often remain persistently affected by neurological deficits. While previous studies suggest that brain-heart electrophysiological interactions may predict severity and prognosis after hypoxic brain injury coma, little is known about the brain-heart dynamics at near-death. Gaining insight into these mechanisms is crucial for developing targeted interventions in critical conditions.

Using a rodent model of reversible systemic anoxia (n=29, male and female rats), we investigated whether brain-heart interactions during the asphyxia onset could predict the return of brain electrical activities after resuscitation. Electrophysiological recordings confirmed that cerebral activity declines following asphyxia, coinciding with increased heart rate variability. Notably, the strong coupling between cardiac parasympathetic activity and high-frequency brain activity in the somatosensory cortex and hippocampus serves as a key predictor of a survival. Our study underscores the potential involvement of the brain-heart axis in the physiology of dying and the potential prognostic significance of the underlying mechanisms, paving the way for translational research into critical care, based on new characterizations of cardiac reflexes and brain-heart interactions.

**Significance Statement:** Understanding the physiological processes that determine survival and recovery following systemic anoxia is critical for improving outcomes in critical care. This study reveals that brain-heart dynamics during the onset of systemic anoxia relate to survival after resuscitation. In particular, the coupling between parasympathetic cardiac activity and high-frequency brain signals. Using a reversible anoxia rodent model, we demonstrate that early brain-heart interactions are not merely consequences of anoxia, but active markers of resilience. These findings offer a novel framework for understanding the physiology of dying and to ultimately developing prognostic tools based on real-time physiological monitoring.

## 1 Introduction

Could electrophysiological changes forecast whether a brain survives or succumbs to a prolonged anoxia? Recent investigations on animal models of respiratory or cardiac arrest have described in detail the alterations in brain electrical activity that accompany the interruption and reinstatement of oxygenation. However, the mechanisms that ultimately lead to brain death remain to be elucidated. A deeper understanding of the physiological mechanisms of dying could enable the development of biomarkers that can predict the outcome of resuscitation procedures (Chawla et al., 2009), such as in-hospital cardiac arrest, or to monitor dying patients after withdrawal of life support.

Consistent cortical dynamics are triggered by systemic anoxia, encompassing an early surge in beta-gamma band activities, followed by a global decline in all frequency bands transiently interrupted by a late relative increase in alpha-delta oscillations (Charpier, 2023). The decrease in amplitude and slowing of cortical activity culminates in a complete cessation of synaptic activity, leading to entry into an isoelectric state characterized by a flat electroencephalogram (EEG) (Borjigin et al., 2013; Li et al., 2015; Schramm et al., 2020; Carton-Leclercq et al., 2023). When anoxia persists, this isoelectric state is transiently interrupted by a large slow wave in the EEG, indicating the collective anoxic depolarization of cortical neurons (Schramm et al., 2020). Interestingly, anoxia-induced cortical changes in rodents are accompanied by sequential changes in cardiac activity that precede its cessation after a few minutes (Li et al., 2015). Specifically, a late increase in coherence between brain and cardiac electrical activity has been reported during the near-death period preceding ventricular fibrillation and asystole (Li et al., 2015). In a large human cohort, it was also shown that the strength of brain-heart electrophysiological interactions were related to the severity of post-anoxic brain injury in patients who had suffered cardiac arrest (Hermann et al., 2024). All this evidence suggests that studying brain-heart interactions could provide valuable insights into the physiology of the dying process. However, we still do not know whether changes in brain and heart activity, and their interactions towards the near death, would define which individuals could survive a resuscitation procedure.

Using a rodent model of reversible systemic anoxia (Schramm et al., 2020; Carton-Leclercq et al., 2023), the present study explores the functional relationship between cardiac functional activity and brain responses to asphyxia to determine whether the dynamics of brain–heart interaction could provide a predictor for survival after a resuscitation procedure. We focused on brain regions - posterior parietal association cortex (PtA), hippocampus (Hpc), thalamus (Th), and primary somatosensory cortex (S1) - that are critically, and potentially differently, affected by hypoxia or anoxia (Kawasaki et al., 1990; Joshi and Andrew, 2001; Tekkök and Ransom, 2004; Schiff, 2008; Macri et al., 2010; Bogdanov et al., 2016). Noteworthy, the analysis of cardiac activity focused on a time-resolved depiction of heart rate variability (HRV), aiming to distinguish the potential contributions of the sympathetic and parasympathetic systems, namely cardiac sympathetic index (CSI) and cardiac parasympathetic index (CPI) (Candia-Rivera et al., 2025).

We found consistent evidence that the progressive decline of brain beta-gamma activity and increase of cardiac responses were correlated with survival. These dynamics were independent of the residual amplitude of brain or heart signals and instead reflected the degree of coordination between brain decay and the emergence of cardiac parasympathetic activity. This suggests that the state of the brain–heart axis in critical conditions may determine resilience and ultimately influence mortality risk.

## 2 Materials and methods

### 2.1 Animals

All procedures were performed in accordance with the European Union Directive 2010/63/EU and received approvals from the national Ministry of Research (APAFIS No.18003-201905101901) and the *Charles Darwin* Ethical Committee (C2EA-05). Experiments were conducted on adult Sprague Dawley rats (male and female, 7 weeks old; Charles River). Rats were kept on a 12-h/12-h light/dark cycle at an average temperature of 25°C and an average humidity of 50%, with ad libitum food and water. Every precaution was taken to minimize stress and number of rodents used in this study.

### 2.2 Surgery and *in vivo* LFP recordings

Surgery and procedures for obtaining local field potentials (LFPs) recordings were essentially performed as previously described (Carton-Leclercq et al., 2023). Briefly, rats (n=29) were first anaesthetized by inhalation of isoflurane (3.5%) and underwent tracheotomy for artificial ventilation (room air, 80 breaths/min, 2.6–3.3 ml/cycle) under neuromuscular blockade (gallamine triethiodide, 40 mg/2h, i.m.). Two small craniotomies were made above the left forelimb region of the S1 (1 mm posterior to bregma and 4–5 mm lateral to the midline)(Paxinos and Watson, 2006) and the left PtA (4.6 mm posterior to bregma and 3.2 mm lateral to the midline). Incisions and pressure points were repeatedly (every 2 h) infiltrated with the local anesthetic agent lidocaine (2%; Centravet). After completion of surgery, isoflurane was discontinued but sedation and analgesia were maintained by repeated injection of sufentanil (3 µg/kg every 30 min, i.p.). This anesthetic condition ensured a globally stable background activity characterized by low amplitude cortical patterns resembling those encountered during waking (Mahon et al., 2001; Bruno and Sakmann, 2006; Altwegg-Boussac et al., 2017). The heart was recorded using bipolar electrocardiographic (ECG) electrodes inserted subcutaneously in the axillary regions. ECG and LFPs were continuously recorded to assess the depth of sedation. Oxygen saturation (SpO_2_), core temperature (maintained at 37°C by a feedback-controlled heating blanket) and end-tidal CO_2_ (EtCO_2_) were also monitored throughout the recording sessions with a veterinary vital signs monitor (LifeWindow unit, Digicare).

Linear 32-channel (200 µm electrode separation distance, 35 µm site IrOx, 0.2 MΩ) silicon probes (ATLAS Neuroengineering®) were inserted perpendicular to the tangent of the S1 surface and perpendicular to the PtA surface to target the PtA, the CA3 region of the Hpc, and the ventro-posteromedial nucleus of the Th (Paxinos and Watson, 2006). Penetrations into the brain tissue were used as the zero reference to determine the depth of recording sites. LFP signals were amplified using a DigitalLynx amplifier (NeuraLynx®), filtered between 0.1 Hz and 6 kHz, and digitized at 32 kHz.

All recordings were visually inspected to identify potential artifacts and to ensure that the signals reflected genuine physiological activity rather than instrumental noise. This quality control was performed by experienced electrophysiologists (authors S.C.-F., S.C., and S.M.). The final cohorts analyzed per region were as follows: PtA, n = 25 (13 deceased, 12 surviving); Hpc, n = 26 (14 deceased, 12 surviving); Th, n = 25 (14 deceased, 11 surviving); and S1, n = 25 (13 deceased, 12 surviving).

### 2.3 Anoxia and rescue protocols

Physiological parameters and LFP activities were continuously monitored during control periods (normoxia), systemic anoxia (asphyxia) and after the restoration of air supply (hereafter referred to as “resuscitation,” denoting only the reoxygenation phase without pharmacological or manual thoracic intervention). Anoxia was induced by a sudden interruption of artificial ventilation in curarized rodents (vent. off). This resulted in an instantaneous collapse of EtCO_2_, followed by a more progressive drop in SpO_2_, reaching undetectable values in less than a minute (Carton-Leclercq et al., 2023; Grou-Radenez et al., 2026).

Resuscitation in this experimental context consisted solely of reinitiating mechanical ventilation (vent. on) to restore cerebral oxygenation and assess spontaneous recovery of brain activity, distinct from the multi-step clinical resuscitation process aimed at cardiovascular stabilization. The duration of anoxia was 187 ± 16.5 s (n = 29), which corresponded to the time at which all structures had undergone anoxic depolarization, a LFP wave marking the onset of the cell death process (Schramm et al., 2020; Charpier, 2023; Grou-Radenez et al., 2026) (see Fig. 3b). At the end of recordings, animals were euthanized with an overdose of euthasol (40%, i.p.) and perfused for subsequent histological processing.

### 2.4 Processing of electrophysiological signals

LFP activities were analyzed using MATLAB R2022b and Fieldtrip Toolbox (Oostenveld et al., 2011). After bandpass (0.5-50 Hz) filtering, fast Fourier transforms were applied to LFP signals to compute their frequency content using a Hanning taper decomposition analysis with a sliding time window of 2 s with a 50% overlap. Frequency-band definitions were established empirically by inspecting group-average time–frequency maps to identify the dominant spectral ranges showing consistent macro-level modulations across the two groups. Three broad bands were thus selected: delta–alpha (1–15 Hz), beta (16–30 Hz), and gamma (31–45 Hz). Such broad grouping, particularly the merged delta–alpha range, is consistent with previous work in comatose and post-anoxic human patients, where low-frequency components jointly reflect cortical suppression due to brain injury and their relationship with outcomes (Hermann et al., 2024, 2025). As a control measure, the delta range (1–4 Hz) was analyzed separately, according to its previous implication in indexing brain damage severity and unconsciousness (Frohlich et al., 2021; Massimini et al., 2024).

ECG time series were bandpass filtered using a Butterworth filter between 0.5 and 50 Hz. The R-peaks from the cardiac cycle were first identified *via* an automatized process, followed by visual inspection and correction of misdetections. The procedure was based on a R-peak template-based cross-correlation and consecutive prominent peaks detection. All the detected peaks were visually inspected over the original ECG, along with the derivative signal of RR series and inter-beat intervals histogram. Manual corrections were performed where needed.

### 2.5 Heart rate and heart rate variability analysis

Cardiac sympathetic and parasympathetic activities were estimated using a recently developed method that exploits the dynamic geometry of the Poincaré plot constructed from inter-beat intervals (IBIs), adapted and validated for rodents (Candia-Rivera et al., 2025). As illustrated in Fig. 1, the analysis begins with the extraction of consecutive IBIs from the ECG signal (Fig. 1a-b). These intervals are used to construct a time-resolved Poincaré plot, in which each IBI is plotted against the subsequent interval (Fig. 1b).

**Figure 1.**
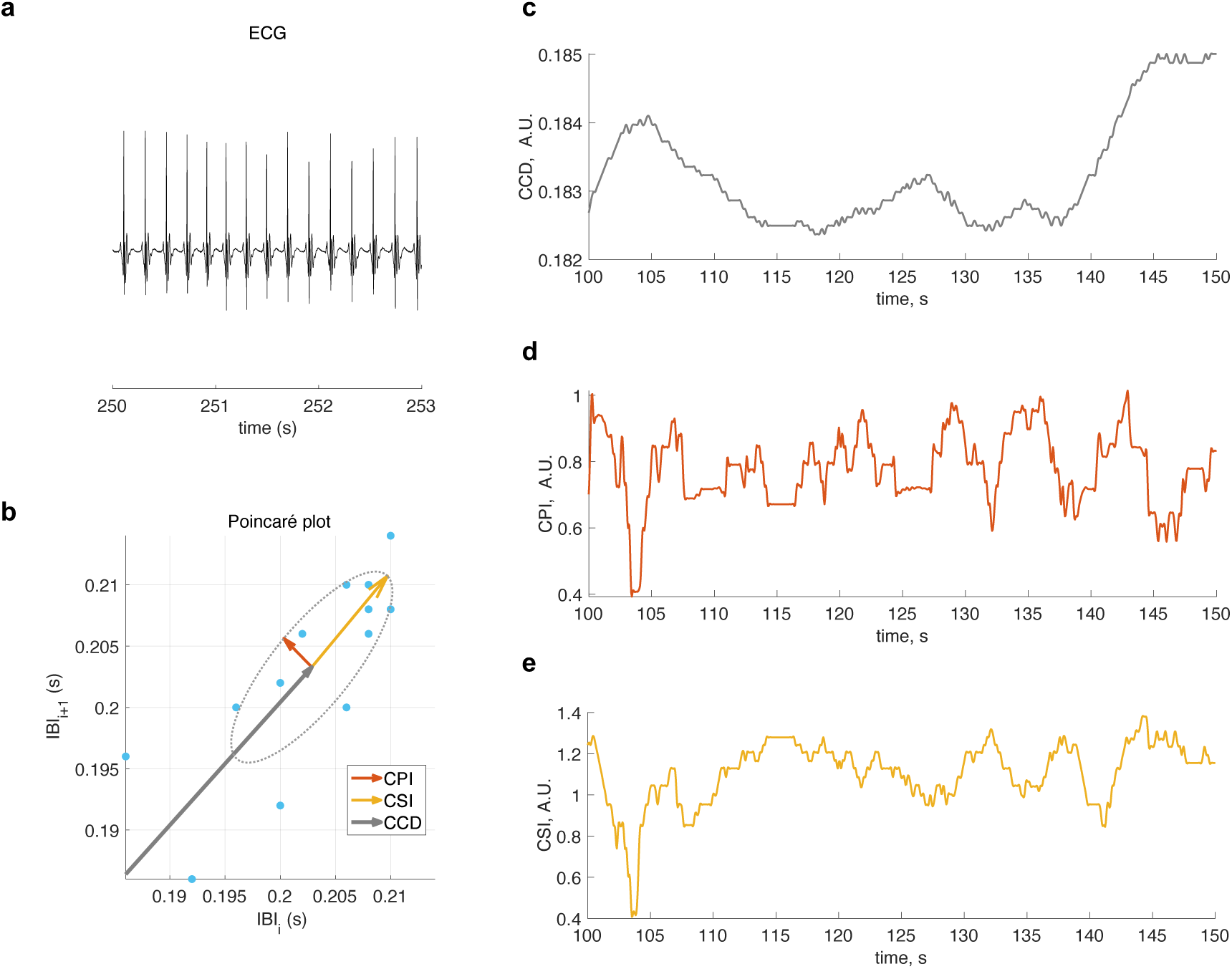
Estimation of time-resolved cardiac autonomic indices from ECG recordings. (a) Representative ECG segment and detection of consecutive inter-beat intervals (IBIs). (b) Poincaré plot constructed from successive IBIs, where each interval IBIi is plotted against the subsequent one, IBIi+1. The ellipse fitted to the point cloud characterizes the geometry of heart rate variability. The short axis of the ellipse (orange) reflects rapid beat-to-beat fluctuations and is used to compute the cardiac parasympathetic index (CPI), whereas the long axis (yellow) reflects slower variability components and is used to compute the cardiac sympathetic index (CSI). (c–e) Time-resolved estimates obtained using a sliding-window approach: (c) cardiac cycle duration (CCD), (d) cardiac parasympathetic index (CPI), and (e) cardiac sympathetic index (CSI). These measures provide complementary information on the temporal dynamics of autonomic cardiac regulation.

The method provides three complementary measures describing cardiac dynamics over time: (i) the cardiac cycle duration (CCD), reflecting beat-to-beat changes in heart period; (ii) the cardiac parasympathetic index (CPI), which quantifies rapid fluctuations of heart rate variability represented by the dispersion of points along the short ratio of the ellipse; and (iii) the cardiac sympathetic index (CSI), which quantifies slower fluctuations represented by the dispersion of points along the long ratio of the ellipse. Together, these measures capture distinct temporal components of autonomic cardiac regulation.

To characterize their temporal evolution, CCD, CPI, and CSI were computed within sliding time windows, yielding continuous time-resolved estimates of cardiac autonomic dynamics (Fig. 1c–e). The computation of these indices is described in Equations 1–3.:

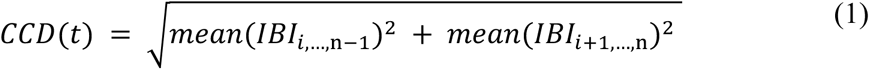

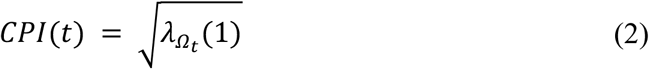

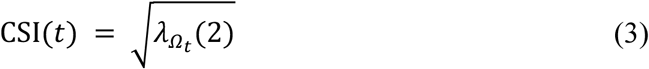

where *λ*_***Ω**t*_(*i*) denotes the i^th^ eigenvalues of a 2 x 2 covariance matrix of *IBI*_i,…,n–1_ and *IBI*_i+1,…,n_, with *Ω*_*t*_: *t* − *T* ≤ *t*_*i*_ ≤ *t*, and *n* is the length of IBI in the time window *Ω*_*t*_. In this study T is set to 3 s, as in previous simulations (Candia-Rivera et al., 2025). To evaluate the robustness of the 3 s window used to compute the cardiac indices, we first identified the latency corresponding to the maximum discrimination between the positive- and negative-outcome groups. We then repeated the discrimination analysis across a range of window lengths (1–15 s) to determine whether the observed group differences were sensitive to the choice of this parameter.

### 2.6 Brain-heart coupling estimation

Brain-heart coupling was assessed by the temporal relationship between fluctuations of LFP power across different frequency bands and the cardiac autonomic indices (Fig. 2). To focus on slow variations in signals amplitude, LFP power and cardiac index time series were transformed into envelope signals. Briefly, local maxima were identified in each time series, and an upper envelope was obtained by spline interpolation between consecutive peaks. The resulting envelope signals corresponded to smoothed LFP power series and cardiac autonomic indices, with reduced high-frequency fluctuations and preserved slower trends associated with autonomic and neural network state changes.

**Figure 2.**
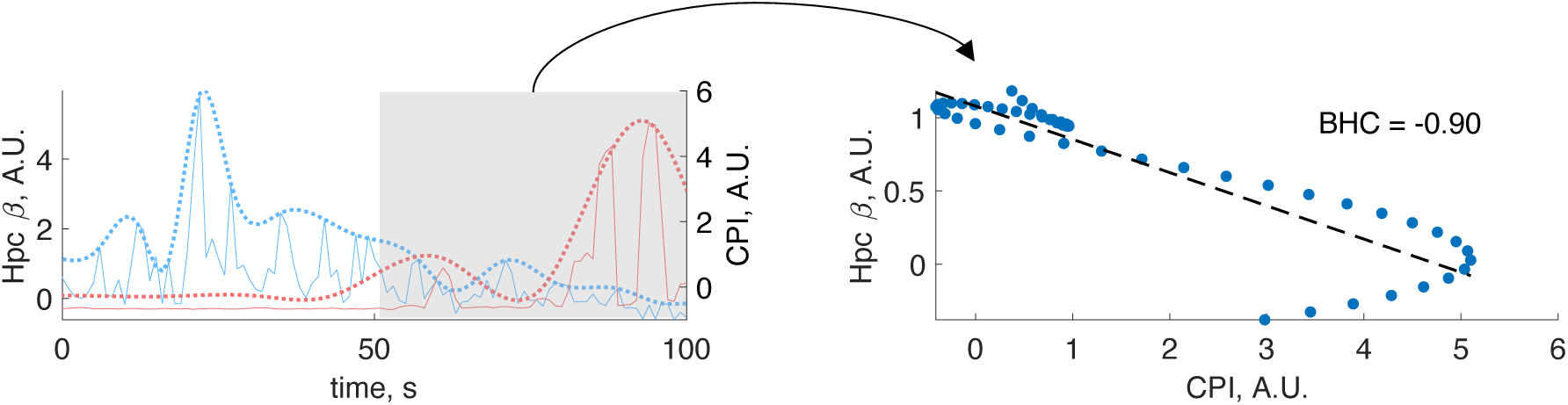
Estimation of brain-heart coupling (BHC). Time-resolved LFP power and cardiac autonomic index signals were first transformed into smooth envelope representations by detecting local maxima and interpolating between successive peaks. The resulting envelope signals capture slow fluctuations in neural and cardiac activity while reducing higher-frequency variability. Brain-heart coupling was quantified as the Spearman correlation coefficient between the LFP power envelope and the cardiac autonomic index envelope within a 50-s sliding window. The example shown illustrates one case, with a strong negative coupling between hippocampal (Hpc) beta-band (◻) power and the cardiac parasympathetic index in the interval 50–100 s (BHC = −0.90). t=0 corresponds to asphyxia onset.

**Figure 3.**
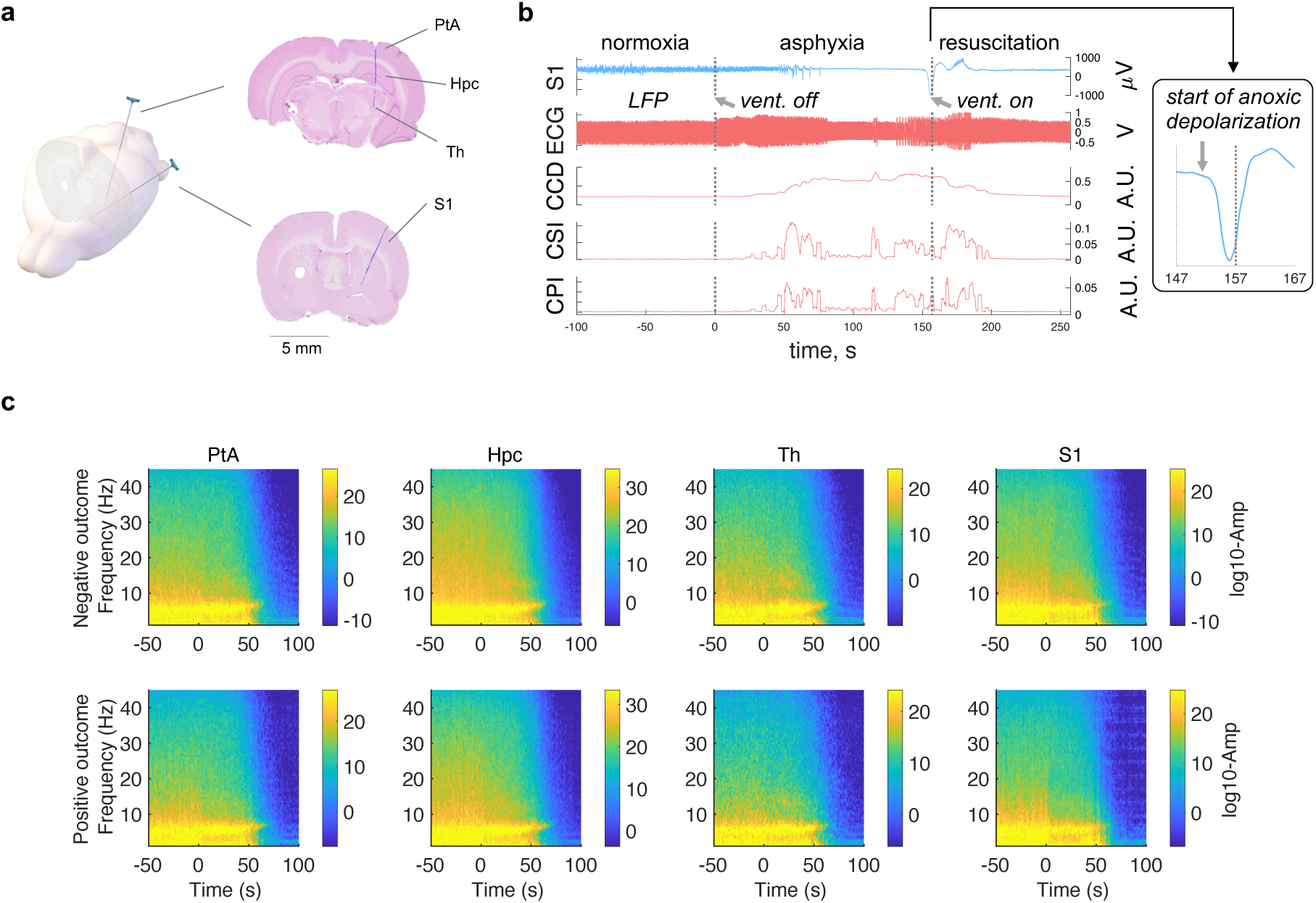
Multi-site LFP and cardiac responses to anoxia onset in the surviving (positive outcome, P.O., n=13) and deceased groups (negative outcome, N.O., n=16)–Part 1. (A) Schematic of the location of recording silicon probes in the parietal association cortex (PtA, n = 25, 13 deceased, 12 surviving), hippocampus (Hpc, n = 26, 14 deceased, 12 surviving), thalamus (Th, n = 25, 14 deceased, 11 surviving), and primary sensory cortex (S1, n = 25, 13 deceased, 12 surviving). See Methods for further details. (B) Typical examples of S1 LFP and ECG records during a control period (normoxia), the period of oxygen deprivation (asphyxia) starting with the interruption of mechanical ventilation (vent. off), and during the rescue process (resuscitation) initiated by the restoration of oxygenation (vent. on). The corresponding changes in the cardiac cycle duration (CCD), the sympathetic (CSI) and parasympathetic (CPI) indices are shown below the traces. The onset at right illustrates the wave of anoxic depolarization (see methods for details). (C) Group average, separated by outcome groups, of the time-frequency changes associated to the asphyxia onset (t=0), in the 1–45 Hz range, and separated by brain region.

Prior to coupling estimation, each envelope time series was normalized to facilitate comparisons across animals and recording sessions. Brain–heart coupling was then quantified using Spearman rank correlation coefficients (Ako et al., 2003). Spearman correlation was selected because it captures monotonic relationships without assuming linearity or normality of the data and is less sensitive to differences in signal amplitude. Consequently, the coupling measure reflects the degree to which neural and cardiac signals co-fluctuate over time rather than their absolute magnitudes.

The temporal window for coupling estimation was determined empirically based on the independent analyses of brain and heart activity performed in successive 50-s intervals. This data-driven definition ensured that the coupling assessment encompassed the period in which both neural and cardiac signals exhibited meaningful changes.

### 2.7 Statistical analysis

Group-wise statistical analysis between the surviving (positive outcome) and deceased (negative outcome) groups was performed through the non-parametric Wilcoxon test for unpaired samples. To control for type-1 errors, we applied permutation test consisting on the resampling of the observations to build an empirical estimate of the null distribution from which the test statistic has been drawn (Kimmel et al., 2007). For each test, p-values were computed as the probability of observing the real data test statistic over the null distribution of test statistics constructed using 10,000 random Monte Carlo permutations. Significance was considered at α<0.05.

Categorical tests were performed though Chi-squared test, in which was compared the proportion of individuals reaching a perfect anticorrelation in the brain-heart coupling measure (correlation coefficient <-0.98) in the positive *vs* negative outcome categories. Significance was considered at α<0.05. Group distributions were reported as group medians.

#Figures displaying box plots indicate the group medians and the box edges indicate the 25^th^ and 75^th^ percentiles.

## 3 Results

We conducted continuous ECG alongside multi-site local LFP recordings in sedated and curarized rats placed under artificial ventilation (n=29) (Fig. 3a and b). Anoxia was induced by interrupting the mechanical ventilation and resuscitation was initiated by the resumption of oxygen supply after 3-4 minutes of anoxia (see Materials and Methods). Once brain oxygenation was restored, 45% of the rats (13 out of 29) gradually regained cortical activities comparable to the normoxic situation (surviving, or positive outcome group), while the others died of cardiac arrest (systemic death) within about 10 minutes (deceased, or negative outcome group).

Four main brain regions were targeted simultaneously: the S1, the PtA, the CA3 region of the Hpc, and the ventro-posteromedial nucleus of the Th (see Fig. 3a). The interruption of artificial ventilation led to a decline in LFP activity across all recorded brain regions, followed by a delayed rebound of low-frequency activity (Fig. 3b). Fig. 3c displays the group-averaged time–frequency representations for the four recorded brain regions (PtA, HPC, Th, and S1) in animals with negative and positive outcomes. These results illustrate the data-driven basis for frequency-band definition, as spectral changes occurred within broad ranges, mostly the 1–15 Hz range, corresponding to the merged delta–alpha band. A gradual reduction in the power of all frequency bands was observed during asphyxia across all regions.

The group-wise depiction of beta-band activity decay in all brain regions is shown in Fig. 4a. S1 was the only region exhibiting marginal differences between the surviving and deceased groups (Fig. 4b). For a detailed visualization of changes in delta-alpha, beta, and gamma ranges, separated by groups, refer to Supplementary Figs. S3-S5,S11). Notably, the low frequencies (delta–alpha range, or delta separately) did not show any discrimination power (Fig. S11).

**Figure 4.**
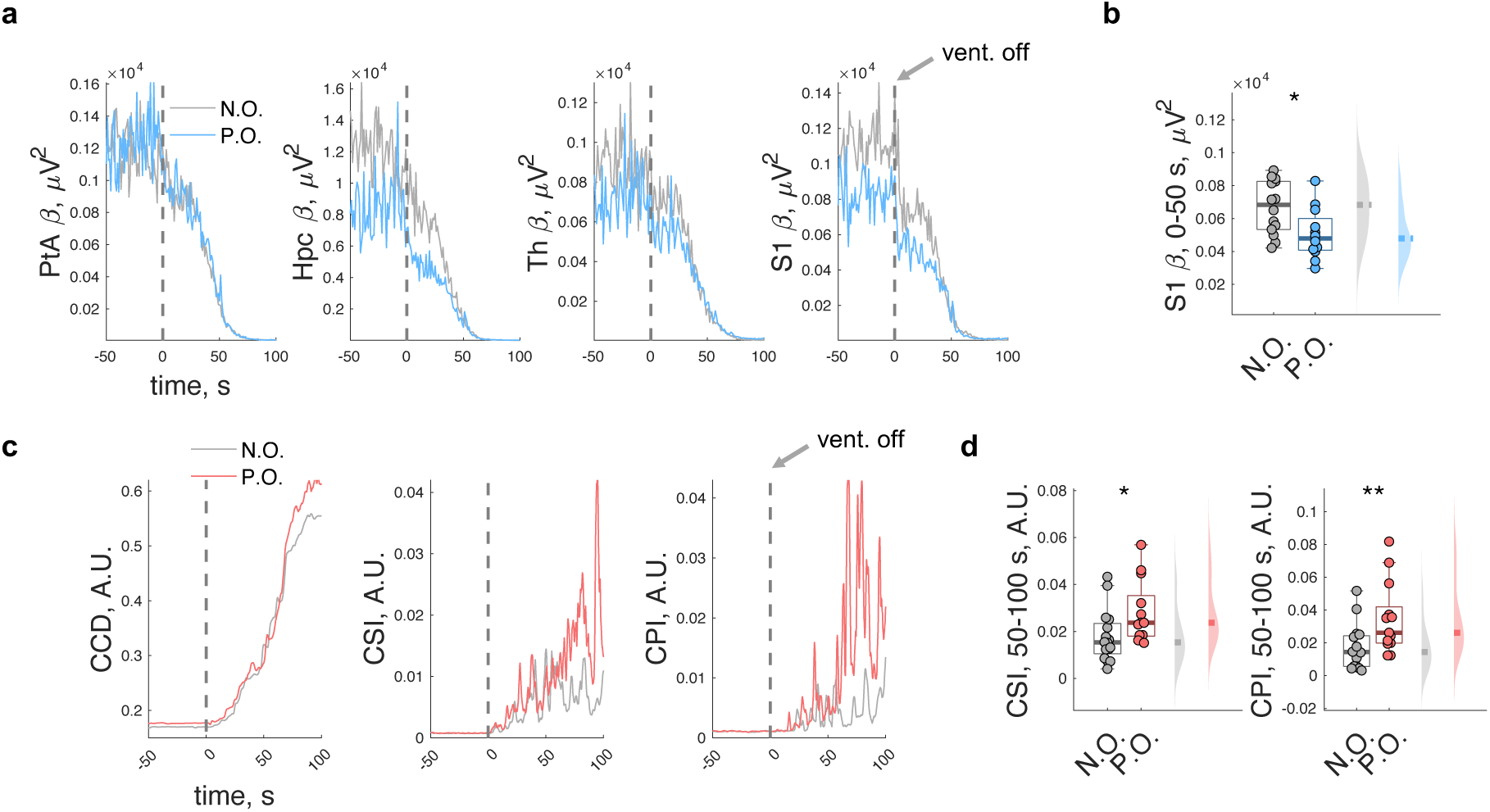
Multi-site LFP and cardiac responses to anoxia onset in the surviving (positive outcome, P.O., n=13) and deceased groups (negative outcome, N.O., n=16)–Part 2. The location of recording silicon probes were: in the parietal association cortex (PtA, n = 25, 13 deceased, 12 surviving), hippocampus (Hpc, n = 26, 14 deceased, 12 surviving), thalamus (Th, n = 25, 14 deceased, 11 surviving), and primary sensory cortex (S1, n = 25, 13 deceased, 12 surviving). (A) Graphs illustrating, for the P.O. group (blue lines) and the N.O. group (grey lines) groups, the evolution of beta activities power in the different brain regions in the 100 s period after vent. off (t=0). (B) In the P.O. group, the decrease in beta activities after vent. off was more pronounced than in the N.O. group (Wilcoxon test, p=0.0119, Z=-2.3917). (C) Graphs showing the evolution of CCD, CSI and CPI in the 100 s period following vent. off. (D) Summary graph of CSI and CPI values in the 50-100 s period after vent. off illustrating that the increase in slow (CSI; Wilcoxon test, p=0.0114, Z=2.4338) and fast (CPI; Wilcoxon test, p=0.0094, Z=2.4777) fluctuations of heart rate variability was more pronounced in the P.O. group.

The gradual collapse of brain electrical activities was accompanied by an increase in HRV, which was particularly pronounced in surviving rodents in both, cardiac sympathetic and parasympathetic indices (Fig. 3c,d; Supplementary Fig. S1). The discriminative power of the cardiac indices remained largely unchanged across sliding-window lengths of 3 s and above, indicating that the results were robust to the choice of this parameter (Supplementary Fig. S15).

We further investigated brain-heart interaction by looking at the degree of correlation between the reduction of brain activity and the increase in HRV. We quantified the strength of brain-heart interactions by computing correlation coefficients between LFP frequency bands and cardiac activity during the 100 s period after asphyxia onset. This window length was defined upon the results showing significant results in LFP power in 0–50 s and in cardiac indices in the 50–100s intervals. The comparison of Spearman correlation coefficients between surviving and deceased groups demonstrated a higher level of coordination between brain and cardiac parasympathetic activities in the group of surviving rodents. This strong negative correlation between brain and cardiac parasympathetic activities in surviving rodents was observed from all brain regions, for both beta and gamma frequency bands, except for gamma activity in the thalamus (Fig. 5; Table 1).

**Figure 5.**
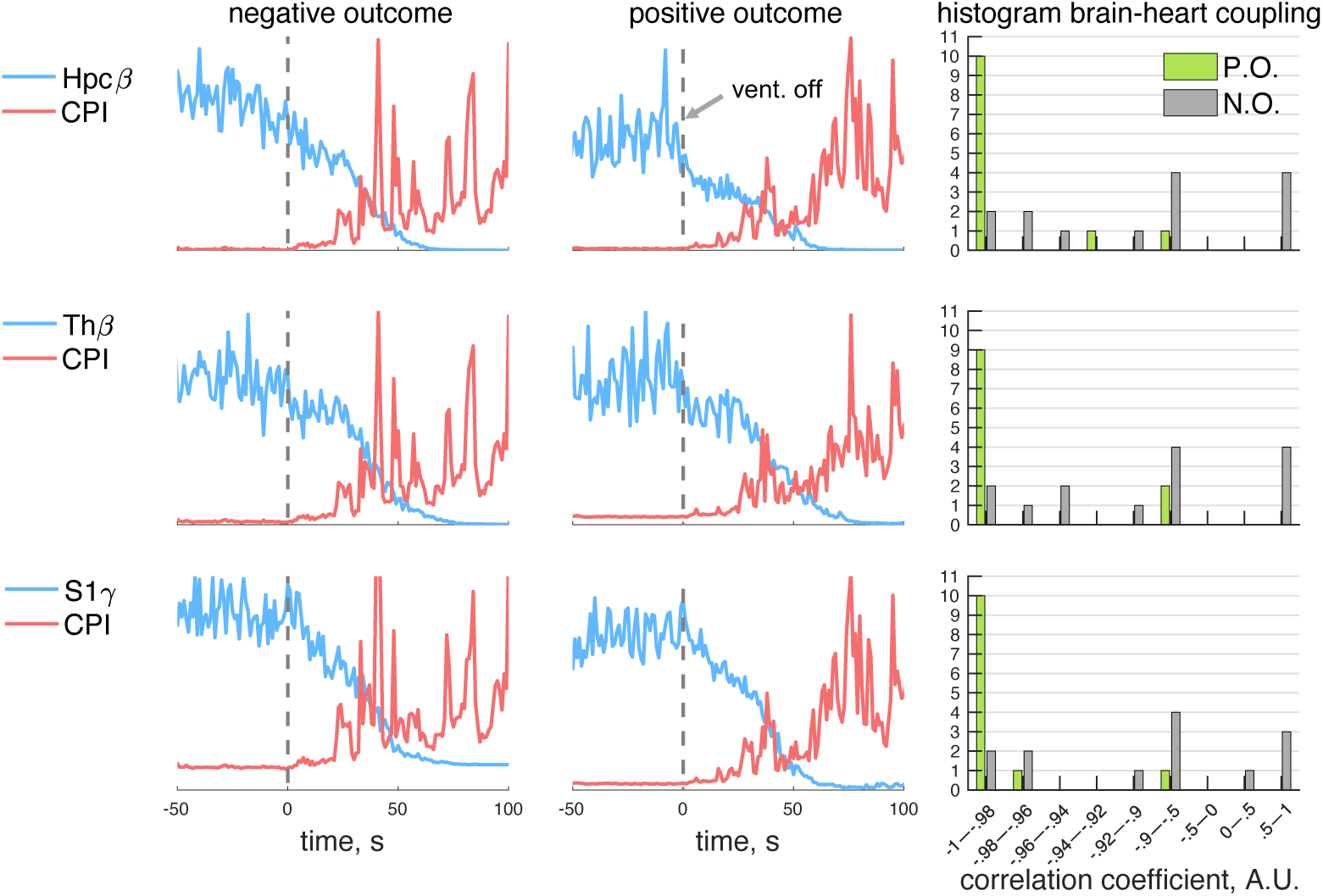
Correlation between multi-site LFP and cardiac parasympathetic responses to asphyxia in the surviving (positive outcome, P.O.) and deceased groups (negative outcome, N.O.). Results are displayed for the hippocampus (Hpc, n = 26, 14 deceased, 12 surviving), thalamus (Th, n = 25, 14 deceased, 11 surviving), and primary sensory cortex (S1, n = 25, 13 deceased, 12 surviving). Group median, z-score normalized, changes in LFP beta (Hpc and Th) or gamma (S1) power (blue lines), together with the temporal evolution of cardiac parasympathetic index (CPI, red lines), in the 100 s period after asphyxia onset (vent. off, t=0). The left graphs correspond to the group of deceased rodents and the middle graphs to the group of surviving rodents. Right, distribution of correlation coefficients quantifying the correlation between LFP power and CPI changes in the 0-50 s time window after anoxia onset in the two groups of rodents. Note that in most individuals with positive outcomes, the negative correlation between brain and cardiac signals was much higher (Spearman correlation coefficient <-0.98).

**Table 1.** Statistical tests performed on the brain-heart coupling estimations. The couplings were computed as the Spearman correlation coefficients between the envelope of beta or gamma power and cardiac series, within 0-50s with respect to the interruption of mechanical ventilation (vent. off). Wilcoxon tests were performed comparing the couplings in surviving vs deceased groups. Chi-square tests were performed comparing the proportion of individuals in each category (surviving vs deceased groups) that presented a correlation coefficient < -0.98.

| Brain region | δ-α LFP – Cardiac fast HRV (CPI) | | $\beta$ LFP – Cardiac fast HRV (CPI) | | $\gamma$ LFP – Cardiac fast HRV (CPI) | |
| --- | --- | --- | --- | --- | --- | --- |
|  | Wilcoxon test | Chi-Square test | Wilcoxon test | Chi-Square test | Wilcoxon test | Chi-Square test |
| PtA | p=0.6438<br>Z=-0.4623 | p=-<br>$\chi^2$=- | **p=0.0036**<br>**Z=-2.8103** | p=9.0771<br>$\chi^2$=0.0026 | **p=0.0367**<br>**Z=-2.0536** | p=0.0254<br>$\chi^2$=4.9958 |
| Hpc | p=0.2076<br>Z=-1.2601 | p=0.0467<br>$\chi^2$= 3.9565 | **p=0.0002**<br>**Z=-3.3923** | **p=0.0004**<br>**$\chi^2$= 12.3957** | **p=0.0021**<br>**Z=-2.8551** | **p=0.0026**<br>**$\chi^2$= 9.1000** |
| Th | p=0.4599<br>Z=-0.7391 | p=0.0962<br>$\chi^2$= 2.7668 | **p=0.0065**<br>**Z=-2.7204** | **p= 0.0007**<br>**$\chi^2$= 11.4016** | p=0.1060<br>Z=-1.6162 | p=0.0868<br>$\chi^2$= 2.9322 |
| S1 | p=0.8918<br>Z=0.1360 | p=-<br>$\chi^2$=- | **p=0.0097**<br>**Z=-2.4844** | **p=0.0021**<br>**$\chi^2$= 9.4195** | **p=0.0021**<br>**Z=-2.8869** | **p=0.0007**<br>**$\chi^2$= 11.5426** |
| Brain region | δ-α LFP – Cardiac slow HRV (CSI) | | $\beta$ LFP – Cardiac slow HRV (CSI) | | $\gamma$ LFP – Cardiac slow HRV (CSI) | |
|  | Wilcoxon test | Chi-Square test | Wilcoxon test | Chi-Square test | Wilcoxon test | Chi-Square test |
| PtA | p=0.4303<br>Z=-0.7887 | p=-<br>$\chi^2$=- | p=0.2216<br>Z=-1.2223 | p=0.1585<br>$\chi^2$=1.9888 | p=0.7015<br>Z=-0.3833 | p=0.3268<br>$\chi^2$=0.9615 |
| Hpc | p=0.8170<br>Z=0.2315 | p=0.1119<br>$\chi^2$=2.5278 | **p=0.0167**<br>**Z=-2.2848** | p=0.0183<br>$\chi^2$=5.5714 | p=0.0945<br>Z=-1.6719 | p=0.0373<br>$\chi^2$=4.3385 |
| Th | p=0.4599<br>Z=0.7391 | p=0.2496<br>$\chi^2$=1.3258 | p=0.2341<br>Z=-1.1899 | p=0.0796<br>$\chi^2$=3.0739 | p=0.7209<br>Z=-0.3572 | p=0.5615<br>$\chi^2$=0.3372 |
| S1 | p=0.6053<br>Z=-0.5167 | p=-<br>$\chi^2$=- | p=0.3110<br>Z=-1.0131 | p=0.0254<br>$\chi^2$=4.9958 | p=0.0768<br>Z=-1.7694 | p=0.0270<br>$\chi^2$=4.8909 |

Bold indicates significance, defined in Wilcoxon test as $p < 0.05$ (Monte Carlo $p$ ) and in Chi-square test as $p < 0.0125$ (Bonferroni corrected $p$ ). PtA: parietal association cortex, Hpc: hippocampus, Th: thalamus, S1: primary somatosensory cortex. “-” indicates that the Chi-square test was not performed because one of the categories had size=0.

Finally, in the period preceding the restoration of brain oxygenation (-100 to -50 seconds), we observed residual activity in some brain structures of the surviving group, which was not seen in the deceased group (see Supplementary Fig. S6-S8). This could result from persistent neuronal activity or could be instrumental noise, as spontaneous brain activity is expected to disappear once the isoelectric state has begun (Schramm et al., 2020; Carton-Leclercq et al., 2023). The resumption of brain oxygenation led to a rapid (0-50 s) regain of LFP activities, which was significantly higher in the surviving group for the delta-alpha range in PtA, Th and S1 (see Supplementary Fig. S8). However, no significant differences between groups for the beta-gamma bands were observed during this period (see Supplementary Fig. S6-S7). Oxygen resumption was also associated with a shorter cardiac cycle duration in the surviving group (see Supplementary Fig. S2), indicative of a sudden increase of the heart rate. With respect to the brain-heart correlations after resuscitation, we did not find significant differences between the two groups (see Supplementary Table S1). The recovery of oxygen was associated with a progressive (after approximately 30 s) decrease in HRV and an increase LFP power in the surviving rodents (see Supplementary Fig. S9).

## 4 Discussion

Prolonged cerebral anoxia following cardiac or respiratory defect is a major cause of death. Following resuscitation, most patients (∼60%) retain permanent neurological deficits of varying severity, mainly due to ischemic or anoxic cortical lesions (Cronberg et al., 2020; Rousseau et al., 2021). Therefore, understanding the physiological mechanisms subtending ongoing cerebral anoxia, degree of resilience and the dying process, remain as important challenges in neuroscience and clinical neurophysiology. This research direction is also relevant for ruling out the physiological mechanisms of the actual process of dying from those associated with near-death experiences (A Shaw, 2024; Martial et al., 2025), which are reported by up to 20% of patients who survive in-hospital cardiac arrest (Mashour et al., 2024).

Recent works on the brain–heart axis have emphasized the integrative nature of neural, mechanical, and biochemical pathways underlying the communication between cerebral, vascular and cardiac dynamics (Valenza et al., 2025; Haque and Dutta, 2026; Scheitz et al., 2026). Such frameworks provide the rationale for translational applications aimed at identifying new prognostic, diagnostic, and therapeutic opportunities derived from brain–heart interactions (Candia-Rivera et al., 2026).

The use of rodent models has recently led to major advances in our understanding of the changes in cortical activity that occur during the cessation and restoration of cerebral oxygenation. Recent studies carried out in animal models of cerebral anoxia (Borjigin et al., 2013; Schramm et al., 2020; Carton-Leclercq et al., 2023), and in patients undergoing withdrawal of end-of-life care (Chawla et al., 2009; Vicente et al., 2022; Xu et al., 2023), have notably shown the early onset of bursts of high-frequency activity (in the beta-gamma band) followed by an overall reduction in electrical activity reflecting a suspension of neuronal and synaptic activity in the cerebral cortex (Schramm et al., 2020; Carton-Leclercq et al., 2023). Our study confirms these findings and shows that the reduction in cerebral electrical activity is linked to an increase in HRV, measured by the cardiac sympathetic and parasympathetic indices. This offers new insights into the potential role of cardiac dynamics in predicting survival after hypoxic or anoxic episodes. Autonomic reflexes, particularly vagal responses (Lemes and Zoccal, 2014), are common following hypoxia and serve as prognostic tools for survival, even during later stages of critical care (Kleiger et al., 1987; Sharshar et al., 2011; Benghanem et al., 2024). Notably, at near death, just before ventricular fibrillation, brain responses to individual heartbeats are significantly reduced (Candia-Rivera and Machado, 2023a). However, occasional strong couplings between brain and cardiac oscillations still occur during this period. These strong couplings have been identified as biomarkers of patient severity and mortality in post-anoxic brain-injured critical care patients (Hermann et al., 2024). Furthermore, during the post-comatose period, brain responses to heartbeats also serve as indicators of consciousness (minimally conscious and cognitive-motor dissociation states) (Candia-Rivera et al., 2021, 2023; Candia-Rivera and Machado, 2023b). Although increased low-frequency brain activity is a well-established hallmark in critical care scenarios (Frohlich et al., 2021; Hermann et al., 2024, 2025; Massimini et al., 2024), emerging from a state of reduced cortical activation and greater neuronal synchronization, which is commonly associated with encephalopathy, disorders of consciousness, brain injury, cerebral ischemia, or sedation. Our analysis revealed that these low-frequency dynamics did not differentiate between positive and negative outcomes when measured within the −50 to 100 s interval relative to asphyxia onset. This suggests that, under the present experimental conditions, low-frequency power alone may not be a reliable predictor of survival.

Our findings therefore expand these previous observations by demonstrating, in a controlled experimental model, how coordinated brain–heart dynamics evolve from asphyxia towards cerebral anoxia. This approach provides a translational bridge between preclinical and human studies, supporting the idea that brain–heart coupling might serve as an integrative biomarker of systemic resilience and physiological state (Dimitri et al., 2022; Podell et al., 2022).

Our findings align with these observations, by emphasizing the role of brain-heart interplay, from the asphyxia stage through to the comatose and post-comatose states. However, the mechanisms we observed still need to be linked to other reports in rodents, including the sequential cardiac responses that precede brain silence (Li et al., 2015), and to findings in humans showing a variable, yet uncoordinated, cessation of brain-heart activity at death (Norton et al., 2017). Additionally, the late increase in coherence between brain and cardiac electrical activity observed during the near-death period (Li et al., 2015), preceding ventricular fibrillation and asystole, may also be relevant to these functional couplings, but further investigation is needed to confirm these links.

By situating these findings within the broader framework of the brain–heart axis, this study contributes to ongoing efforts to characterize how integrative physiological processes breakdown or reorganize during critical transitions, with potential implications for monitoring and intervention in neurocritical and cardiovascular care (Shi et al., 2026).

We found a significant relationship between cardiac responses and the S1 during asphyxia, consistent with studies reporting that the S1 is highly responsive to anoxic states. Optical imaging and multi-scale electrophysiological experiments have shown that anoxic depolarization is preferentially initiated in S1, by the pyramidal neurons of the deep layers (Bogdanov et al., 2016; Schramm et al., 2020; Carton-Leclercq et al., 2023). Notably, S1 is also extensively involved in integrating bodily rhythms, including cardiac, respiratory, and gastric activity (Engelen et al., 2023), suggesting that its susceptibility to depolarization may have broad physiological and clinical implications.

Although the hippocampus is well known for its sensitivity to oxygen deprivation and other critical states (Lipton, 1999; Chawla et al., 2009), our results did not identify it as a region most vulnerable to anoxia as compared to the other brain structures investigated here, at least in the time window preceding terminal depolarization. We found a strong link between cardiac dynamics and hippocampus activity. While previous evidence demonstrated an increase in hippocampal activity and heightened connectivity between the motor and prefrontal regions in gamma frequencies during critical physiological states (Zhang et al., 2019; Xu et al., 2023), electrophysiological measures alone often failed to provide meaningful insights into the patient’s physiological state during resuscitation. However, we did detect coupling between hippocampal activity and cardiac dynamics, suggesting that, while the hippocampus may not depolarize first, it remains functionally engaged with autonomic processes until late in the dying sequence. Future studies incorporating multi-scale recordings or selective perfusion mapping could further elucidate regional differences in vulnerability and brain–heart coupling under terminal hypoxia.

On the heart side, the role of sudden HRV in critical contexts, including hypoxia, remained unknown (Zou et al., 2017). Our results suggest a coordinated physiological response between oscillatory activity in different brain regions and cardiac responses at near-death and underlines its significance for predicting mortality.

We cannot completely rule out the presence of confounding factors related to our sedation or curarization procedures, but it is unlikely that they influenced our results. While fentanyl derivatives in high doses can decrease heart rate (Freye, 1974) and gallamine triethiodide can increase heart rate (Brown and Crout, 1970), our main results focus on rhythm fluctuations associated with vagal tone rather than absolute heart rate, indicating that the observed brain-heart interactions are not primarily due to these agents. Similarly, although HRV is often used to assess cardiac outcomes, in our model the observed effects reflect coordinated brain-heart dynamics rather than changes in HRV measures, which have shown limited predictive value in those critical scenarios (Benghanem et al., 2024; Kula et al., 2025).

Our brain–heart coupling analysis acknowledges some limitations. First, the extraction of signal envelopes attenuates fast temporal fluctuations, potentially reducing sensitivity to transient dynamics in the data. This limitation arises because envelope estimation effectively smooths the underlying signal, emphasizing slower oscillatory components. Second, the accuracy of the envelope representation may depend on the performance of the peak detection algorithm, which may vary with signal quality and noise characteristics. It is also possible that the higher-frequency fluctuations attenuated by the envelope carry additional physiological or clinical significance. However, in the present study, the discriminatory power was driven by slow components, aligning with our focus on low-frequency power fluctuations in brain–heart co-fluctuations. Beyond these technical considerations, the proposed framework represents a descriptive, correlation-based approach rather than mechanistic, serving as a simple yet important first step toward linking brain and heart dynamics to the physiology of the dying process. Future studies that integrate complementary data, such as functional outcomes or behavioral measures, may further elucidate the contribution of high-frequency co-fluctuations and help bridge the gap between descriptive observations and mechanistic understanding.

Although our study focused on the physiological dynamics preceding death rather than on long-term functional outcomes, we recognize that variability in the extent and spatial pattern of anoxia-induced brain damage could contribute to inter-individual differences in electrophysiological measures. However, due to the experimental design, we were not able to control on the lesion-specific dynamics. Future studies combining longitudinal recordings with assessment of anoxic/ischemic neuronal injury could help clarify how regional vulnerability and lesion severity relate to specific electrophysiological signatures and recovery trajectories following anoxia.

While EEG remains a valuable tool (Rossetti et al., 2012; Ruijter et al., 2019; Benghanem et al., 2022), our findings add to the growing body of evidence highlighting these electrophysiological markers in preclinical scenarios (Roberti et al., 2023), but also the importance of brainstem-autonomic reflexes, interoceptive mechanisms and brain-heart interactions in prognosis following critical scenarios (Sharshar et al., 2011; Benghanem et al., 2024; Hermann et al., 2024, 2025; Bouchereau et al., 2025). By integrating these physiological systems, future prognostication efforts could become more comprehensive and accurate. These investigations not only shed light on prognosis but also probe fundamental issues concerning the functional role of the brain-heart connection. Although the anoxic-dependent mechanisms remain elusive, brain-heart communication is known to involve multiple pathways, including neurovegetative, endocrine, immune, and genetic regulation mechanisms, as seen in conditions like stroke (Chen et al., 2017).

Understanding these multilevel pathways may ultimately inform the design of strategies that promote brain recovery through interventions such as autonomic modulation, structured breathing, or real-time physiological feedback, aiming to enhance survival and neurological outcome. It is important to note that the present findings do not imply a purely brain-to-heart influence. While our analysis revealed significant co-fluctuations between neural and cardiac activities, the directionality of these interactions was not assessed. Given that the somatosensory cortex is part of the interoceptive network and continuously integrates visceral and autonomic inputs (Engelen et al., 2023), the observed coupling may also reflect heart-to-brain signaling. This bidirectional perspective aligns with recent views of the brain–heart axis as a dynamic feedback system rather than a unidirectional control pathway (Valenza et al., 2025; Candia-Rivera et al., 2026). Future studies incorporating directional connectivity measures will be needed to disentangle ascending and descending contributions to these co-fluctuations during the dying process.

Uncovering these complexities could lead to new methods for analyzing brain-heart interactions and their role in neural reorganization before and after severe neuronal injury. In summary, our findings align with current translational trends emphasizing the brain–heart axis as a key determinant of physiological stability and recovery, underlining the potential clinical utility of multimodal monitoring techniques to assess systemic integrity and prognosis in critical conditions.

## Author contribution statement

Diego Candia-Rivera: Conceptualization, Methodology, Software, Validation, Formal Analysis, Investigation, Writing Original Draft, Writing Review and Editing.

Sofia Carrion-Falgarona: Data Curation, Methodology, Validation, Investigation, Writing Review and Editing.

Mario Chavez: Methodology, Investigation, Supervision, Writing Review and Editing. Fabrizio de Vico Fallani: Methodology, Investigation, Writing Review and Editing.

Stéphane Charpier: Methodology, Investigation, Resources, Supervision, Writing Review and Editing.

Séverine Mahon: Methodology, Validation, Investigation, Resources, Supervision, Writing Review and Editing.

## Acknowledgments

This work was supported by the European Commission, Horizon MSCA Postdoctoral Fellowship Program, to Diego Candia-Rivera (grant n° 101151118).

## References

A Shaw N (2024) The gamma-band activity model of the near-death experience: a critique and a reinterpretation. F1000Res 13:674.

Ako M, Kawara T, Uchida S, Miyazaki S, Nishihara K, Mukai J, Hirao K, Ako J, Okubo Y (2003) Correlation between electroencephalography and heart rate variability during sleep. Psychiatry Clin Neurosci 57:59–65.

Altwegg-Boussac T, Schramm AE, Ballestero J, Grosselin F, Chavez M, Lecas S, Baulac M, Naccache L, Demeret S, Navarro V, Mahon S, Charpier S (2017) Cortical neurons and networks are dormant but fully responsive during isoelectric brain state. Brain 140:2381–2398.

Benghanem S, Pruvost-Robieux E, Bouchereau E, Gavaret M, Cariou A (2022) Prognostication after cardiac arrest: how EEG and evoked potentials may improve the challenge. Ann Intensive Care 12:111.

Benghanem S, Sharshar T, Gavaret M, Dumas F, Diehl J-L, Brechot N, Picard F, Candia-Rivera D, Le M-P, Pène F, Cariou A, Hermann B (2024) Heart Rate Variability for Neuro-Prognostication After CA: Insight from the Parisian Registry. Resuscitation.

Bogdanov VB, Middleton NA, Theriot JJ, Parker PD, Abdullah OM, Ju YS, Hartings JA, Brennan KC (2016) Susceptibility of Primary Sensory Cortex to Spreading Depolarizations. J Neurosci 36:4733–4743.

Borjigin J, Lee U, Liu T, Pal D, Huff S, Klarr D, Sloboda J, Hernandez J, Wang MM, Mashour GA (2013) Surge of neurophysiological coherence and connectivity in the dying brain. Proceedings of the National Academy of Sciences 110:14432–14437.

Bouchereau E, Pruvost-Robieux E, Siami S, Chaffaut C, Bouglé A, Gavaret M, Heming N, Sivanandamoorthy S, Zyss J, Degos V, Kandelman S, Righy Shinotsuka C, Benghanem S, Naccache L, Rohaut B, Hermann B, Azabou E, Chevret S, Sharshar T (2025) Altered lower brainstem neurophysiological response is associated with mortality in deeply sedated critically ill patients. Intensive Care Med 51:1050–1061.

Brown BR, Crout JR (1970) The sympathomimetic effect of gallamine on the heart. J Pharmacol Exp Ther 172:266–273.

Bruno RM, Sakmann B (2006) Cortex is driven by weak but synchronously active thalamocortical synapses. Science 312:1622–1627.

Candia-Rivera D, Annen J, Gosseries O, Martial C, Thibaut A, Laureys S, Tallon-Baudry C (2021) Neural Responses to Heartbeats Detect Residual Signs of Consciousness during Resting State in Postcomatose Patients. J Neurosci 41:5251–5262.

Candia-Rivera D, Carrion-Falgarona S, Fallani F de V, Chavez M (2025) Modeling the time-resolved modulations of cardiac activity in rats: A study on pharmacological autonomic stimulation. Journal of Physiology in press.

Candia-Rivera D, Faes L, Fallani FDV, Chavez M (2026) Measures and Models of Brain-Heart Interactions. IEEE Reviews in Biomedical Engineering 19:24–40.

Candia-Rivera D, Machado C (2023a) Reduced Heartbeat-Evoked Responses in a Near-Death Case Report. Journal of Clinical Neurology 19:581–588.

Candia-Rivera D, Machado C (2023b) Multidimensional assessment of heartbeat-evoked responses in disorders of consciousness. European Journal of Neuroscience 58:3098–3110.

Candia-Rivera D, Raimondo F, Pérez P, Naccache L, Tallon-Baudry C, Sitt JD (2023) Conscious processing of global and local auditory irregularities causes differentiated heartbeat-evoked responses King AJ, Garrido M, Chait M, eds. eLife 12:e75352.

Carton-Leclercq A, Carrion-Falgarona S, Baudin P, Lemaire P, Lecas S, Topilko T, Charpier S, Mahon S (2023) Laminar organization of neocortical activities during systemic anoxia. Neurobiology of Disease 188:106345.

Charpier S (2023) Between life and death: the brain twilight zones. Front Neurosci 17.

Chawla LS, Akst S, Junker C, Jacobs B, Seneff MG (2009) Surges of electroencephalogram activity at the time of death: a case series. J Palliat Med 12:1095–1100.

Chen Z, Venkat P, Seyfried D, Chopp M, Yan T, Chen J (2017) Brain–Heart Interaction. Circulation Research 121:451–468.

Cronberg T, Greer DM, Lilja G, Moulaert V, Swindell P, Rossetti AO (2020) Brain injury after cardiac arrest: from prognostication of comatose patients to rehabilitation. Lancet Neurol 19:611–622.

Dimitri GM, Beqiri E, Placek MM, Czosnyka M, Stocchetti N, Ercole A, Smielewski P, Lió P, CENTER-TBI Collaborators (2022) Modeling Brain-Heart Crosstalk Information in Patients with Traumatic Brain Injury. Neurocrit Care 36:738–750.

Engelen T, Solcà M, Tallon-Baudry C (2023) Interoceptive rhythms in the brain. Nat Neurosci 26:1670–1684.

Freye E (1974) Cardiovascular Effects of High Dosages of Fentanyl, Meperdine, and Naloxone in Dogs. Anesthesia & Analgesia 53:40.

Frohlich J, Toker D, Monti MM (2021) Consciousness among delta waves: a paradox? Brain 144:2257–2277.

Grou-Radenez A, Carton-Leclercq A, Carrion-Falgarona S, Pinto A, Oudea B, Lecas S, Charpier S, Mahon S (2026) Cortical network resilience to transient anoxia involves redistribution of single-neuron firing. Neurobiology of Disease 226:107441.

Haque S, Dutta P (2026) The Heart-Brain Axis in the Context of Cardiovascular Disease. Physiology (Bethesda) 41:0.

Hermann B, Benghanem S, Pruvost-Robieux E, Sharshar T, Gavaret M, Cariou A, Diehl J-L, Candia-Rivera D (2025) Brain-heart interactions are associated with mortality and acute encephalopathy in ICU patients with severe COVID-19. Clinical Neurophysiology 175:2110745.

Hermann B, Candia-Rivera D, Sharshar T, Gavaret M, Diehl J-L, Cariou A, Benghanem S (2024) Aberrant brain–heart coupling is associated with the severity of post cardiac arrest brain injury. Annals of Clinical and Translational Neurology 11:866–882.

Joshi I, Andrew RD (2001) Imaging Anoxic Depolarization During Ischemia-Like Conditions in the Mouse Hemi-Brain Slice. Journal of Neurophysiology 85:414–424.

Kawasaki K, Traynelis SF, Dingledine R (1990) Different responses of CA1 and CA3 regions to hypoxia in rat hippocampal slice. J Neurophysiol 63:385–394.

Kimmel G, Jordan MI, Halperin E, Shamir R, Karp RM (2007) A Randomization Test for Controlling Population Stratification in Whole-Genome Association Studies. Am J Hum Genet 81:895–905.

Kleiger RE, Miller JP, Bigger JT, Moss AJ (1987) Decreased heart rate variability and its association with increased mortality after acute myocardial infarction. Am J Cardiol 59:256–262.

Kula Y, Wacht O, Ben Shlomo I, Gitler A, Gidron Y (2025) Does heart rate variability predict and improve performance in pediatric CPR?-a simulation study. BMC Emerg Med 25:52.

Lemes EV, Zoccal DB (2014) Vagal afferent control of abdominal expiratory activity in response to hypoxia and hypercapnia in rats. Respir Physiol Neurobiol 203:90–97.

Li D, Mabrouk OS, Liu T, Tian F, Xu G, Rengifo S, Choi SJ, Mathur A, Crooks CP, Kennedy RT, Wang MM, Ghanbari H, Borjigin J (2015) Asphyxia-activated corticocardiac signaling accelerates onset of cardiac arrest. Proc Natl Acad Sci U S A 112:E2073–2082.

Lipton P (1999) Ischemic cell death in brain neurons. Physiol Rev 79:1431–1568.

Macri MA, D’Alessandro N, Di Giulio C, Di Iorio P, Di Luzio S, Giuliani P, Esposito E, Pokorski M (2010) Region-specific effects on brain metabolites of hypoxia and hyperoxia overlaid on cerebral ischemia in young and old rats: a quantitative proton magnetic resonance spectroscopy study. Journal of Biomedical Science 17:14.

Mahon S, Deniau J-M, Charpier S (2001) Relationship between EEG Potentials and Intracellular Activity of Striatal and Cortico-striatal Neurons: an In Vivo Study under Different Anesthetics. Cerebral Cortex 11:360–373.

Martial C, Fritz P, Gosseries O, Bonhomme V, Kondziella D, Nelson K, Lejeune N (2025) A neuroscientific model of near-death experiences. Nat Rev Neurol:1–15.

Mashour GA, Lee U, Pal D, Li D (2024) Consciousness and the Dying Brain. Anesthesiology 140:1221–1231.

Massimini M, Corbetta M, Sanchez-Vives MV, Andrillon T, Deco G, Rosanova M, Sarasso S (2024) Sleep-like cortical dynamics during wakefulness and their network effects following brain injury. Nat Commun 15:7207.

Norton L, Gibson RM, Gofton T, Benson C, Dhanani S, Shemie SD, Hornby L, Ward R, Young GB (2017) Electroencephalographic Recordings During Withdrawal of Life-Sustaining Therapy Until 30 Minutes After Declaration of Death. Canadian Journal of Neurological Sciences 44:139–145.

Oostenveld R, Fries P, Maris E, Schoffelen J-M (2011) FieldTrip: Open Source Software for Advanced Analysis of MEG, EEG, and Invasive Electrophysiological Data. Computational Intelligence and Neuroscience 2011:9 pages.

Paxinos G, Watson C (2006) The rat brain in stereotaxic coordinates. Elsevier.

Podell J, Pergakis M, Yang S, Felix R, Parikh G, Chen H, Chen L, Miller C, Hu P, Badjatia N (2022) Leveraging Continuous Vital Sign Measurements for Real-Time Assessment of Autonomic Nervous System Dysfunction After Brain Injury: A Narrative Review of Current and Future Applications. Neurocrit Care 37:206–219.

Roberti E, Chiarini G, Latronico N, Adami EC, Plotti C, Bonetta E, Magri F, Rasulo FA, Coma following Cardiac ArreST study group (COAST) (2023) Electroencephalographic monitoring of brain activity during cardiac arrest: a narrative review. Intensive Care Med Exp 11:4.

Rossetti AO, Carrera E, Oddo M (2012) Early EEG correlates of neuronal injury after brain anoxia. Neurology 78:796–802.

Rousseau A-F, Prescott HC, Brett SJ, Weiss B, Azoulay E, Creteur J, Latronico N, Hough CL, Weber-Carstens S, Vincent J-L, Preiser J-C (2021) Long-term outcomes after critical illness: recent insights. Crit Care 25:108.

Ruijter BJ, Tjepkema-Cloostermans MC, Tromp SC, van den Bergh WM, Foudraine NA, Kornips FHM, Drost G, Scholten E, Bosch FH, Beishuizen A, van Putten MJAM, Hofmeijer J (2019) Early electroencephalography for outcome prediction of postanoxic coma: A prospective cohort study. Annals of Neurology 86:203–214.

Scheitz JF, Villringer A, Mikail N, Gebhard C, Endres M (2026) Bidirectional brain–heart interactions in health and disease. Nat Rev Neurol 22:209–225.

Schiff ND (2008) Central Thalamic Contributions to Arousal Regulation and Neurological Disorders of Consciousness. Annals of the New York Academy of Sciences 1129:105–118.

Schramm AE, Carton-Leclercq A, Diallo S, Navarro V, Chavez M, Mahon S, Charpier S (2020) Identifying neuronal correlates of dying and resuscitation in a model of reversible brain anoxia. Progress in Neurobiology 185:101733.

Sharshar T et al. (2011) Brainstem responses can predict death and delirium in sedated patients in intensive care unit. Crit Care Med 39:1960–1967.

Shi Q, Shi L, Chao J, Liu J, Zhang L, Tian F, Xu C, Hu B (2026) Exploring brain-heart interactions: advances in physiological signal fusion for healthcare. Information Fusion 128:103950.

Tekkök SB, Ransom BR (2004) Anoxia effects on CNS function and survival: regional differences. Neurochem Res 29:2163–2169.

Valenza G, Matić Z, Catrambone V (2025) The brain–heart axis: integrative cooperation of neural, mechanical and biochemical pathways. Nat Rev Cardiol 22:537–550.

Vicente R, Rizzuto M, Sarica C, Yamamoto K, Sadr M, Khajuria T, Fatehi M, Moien-Afshari F, Haw CS, Llinas RR, Lozano AM, Neimat JS, Zemmar A (2022) Enhanced Interplay of Neuronal Coherence and Coupling in the Dying Human Brain. Frontiers in Aging Neuroscience 14 Available at: https://www.frontiersin.org/articles/10.3389/fnagi.2022.813531 [Accessed September 10, 2022].

Xu G, Mihaylova T, Li D, Tian F, Farrehi PM, Parent JM, Mashour GA, Wang MM, Borjigin J (2023) Surge of neurophysiological coupling and connectivity of gamma oscillations in the dying human brain. Proceedings of the National Academy of Sciences 120:e2216268120.

Zhang Y, Li Z, Zhang J, Zhao Z, Zhang H, Vreugdenhil M, Lu C (2019) Near-Death High-Frequency Hyper-Synchronization in the Rat Hippocampus. Front Neurosci 13.

Zou R, Shi W, Tao J, Li H, Lin X, Yang S, Hua P (2017) Neurocardiology: Cardiovascular Changes and Specific Brain Region Infarcts. BioMed Research International 2017:e5646348.

