## Supplementary material for "Brain–heart interactions underlying differential recovery after transient systemic anoxia" (from another version of this preprint: version 1, 2026-04-21)

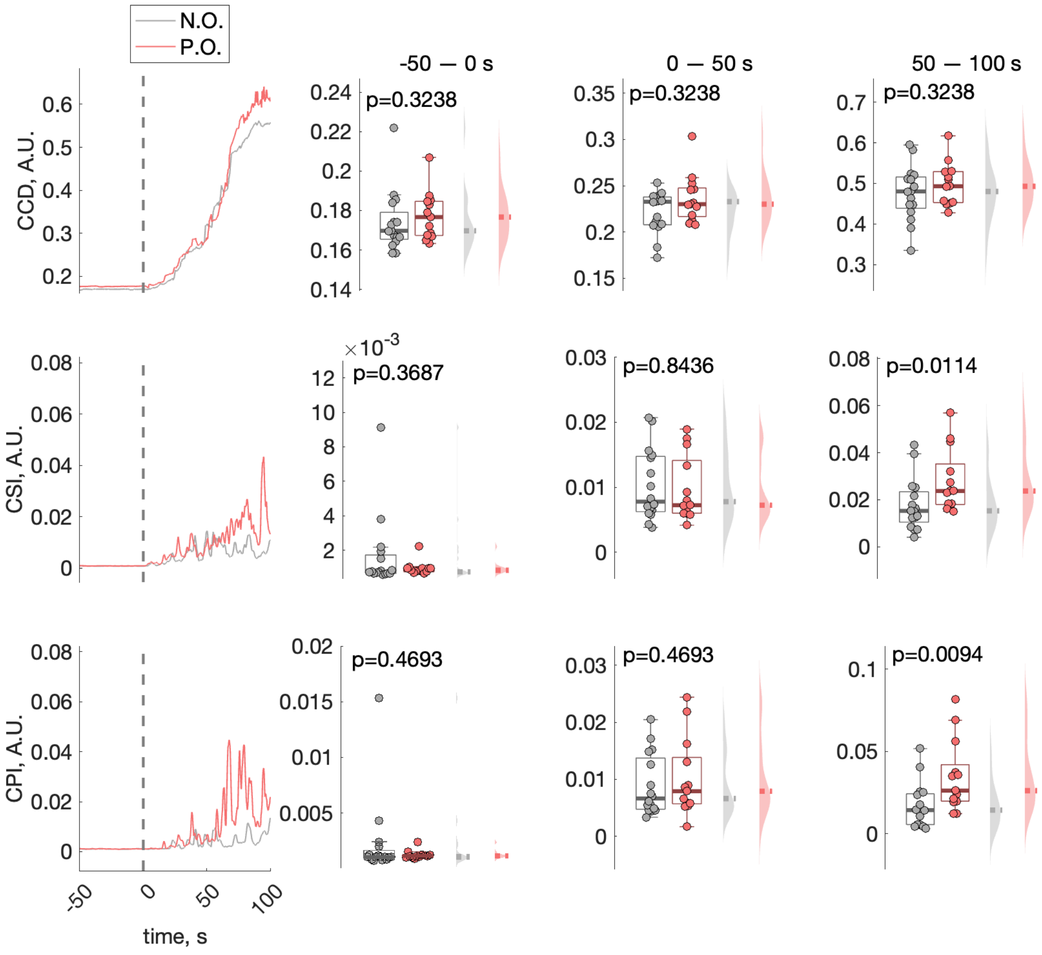


Figure S1. *Cardiac responses to asphyxia. The cardiac cycle duration (CCD), the cardiac sympathetic (CSI) and parasympathetic (CSI) indices were computed before and after the induction of a systemic anoxia (t=0, thick dashed grey line) in rodents recovering brain activities (positive outcome, P.O., red lines) or dying rapidly (negative outcome, N.O., grey lines) after a resuscitation protocol. As shown in the middle and bottom leftmost graphs, CSI and CPI indices are significantly increased 50-100 s after the onset of asphyxia in the P.O. group as compared to the N.O. group. Thick lines in the box plots indicate median value, edges indicate the 25th to 75th percentiles, and whiskers extend up to 1.5 times the interquartile range from the edges.*

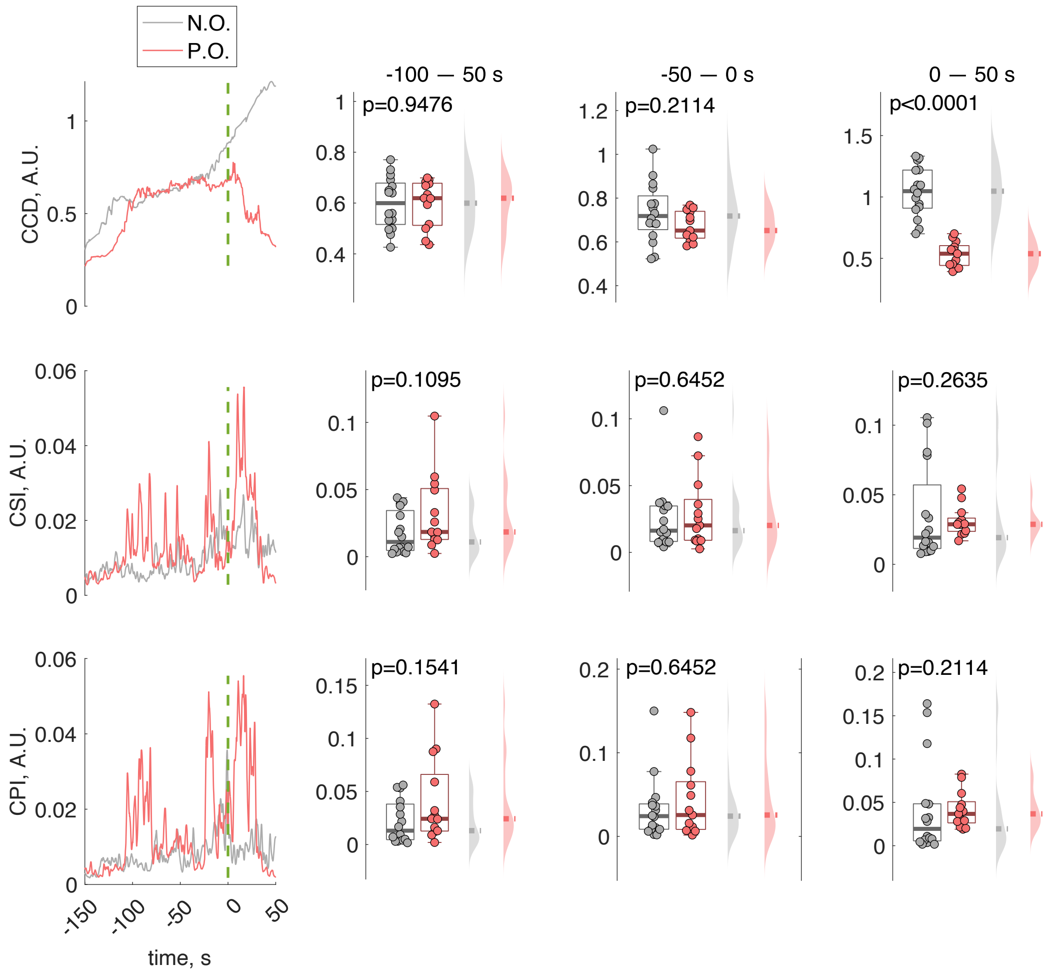


Figure S2. *Cardiac responses to resuscitation. The cardiac cycle duration (CCD), the cardiac sympathetic (CSI) and parasympathetic (CSI) indices were computed before and after the induction of a systemic anoxia (t=0, thick dashed green line) in rodents recovering brain activities (positive outcome, P.O., red lines) or dying rapidly (negative outcome, N.O., grey lines) after a resuscitation protocol. CCD differed between groups at 0-50 s, while CSI and CPI values did not significantly differ between the two groups. Thick lines in the box plots indicate median value, edges indicate the 25th to 75th percentiles, and whiskers extend up to 1.5 times the interquartile range from the edges.*

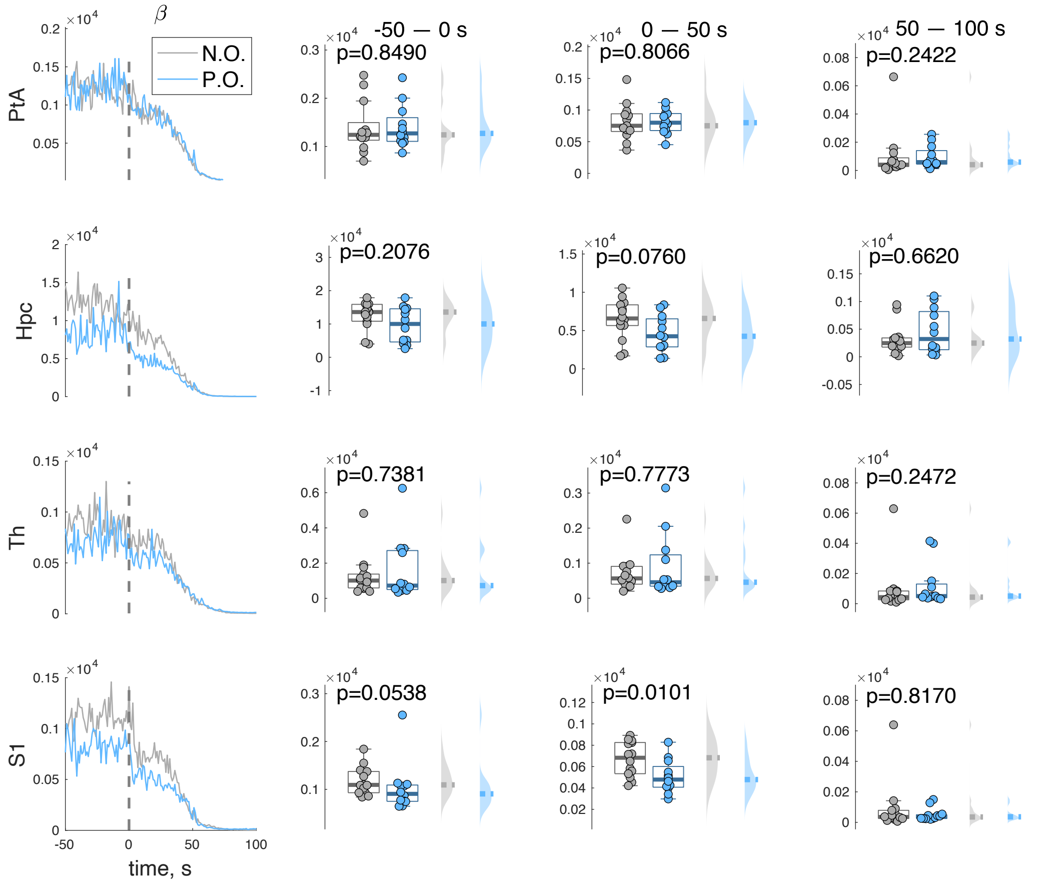


Figure S3. *Changes in the beta band LFP activities following anoxia onset. The power of beta (β) activities in LFP records from the parietal association cortex (PtA), hippocampus (HpC), thalamus (Th), and primary somatosensory cortex (S1) was computed before and after cessation of brain oxygenation (t=0, thick dashed gray line). The decrease in S1 beta activities in the 0-50 s time window after asphyxia was found significantly higher in the group of rodents with positive outcome (P.O., blue signals) than in rodents dying after the resuscitation attempt (N.O., grey signals). Changes in beta activities in the other brain structures did not differ between the two groups. Thick lines in the box plots indicate median value, edges indicate the 25th to 75th percentiles, and whiskers extend up to 1.5 times the interquartile range from the edges.*

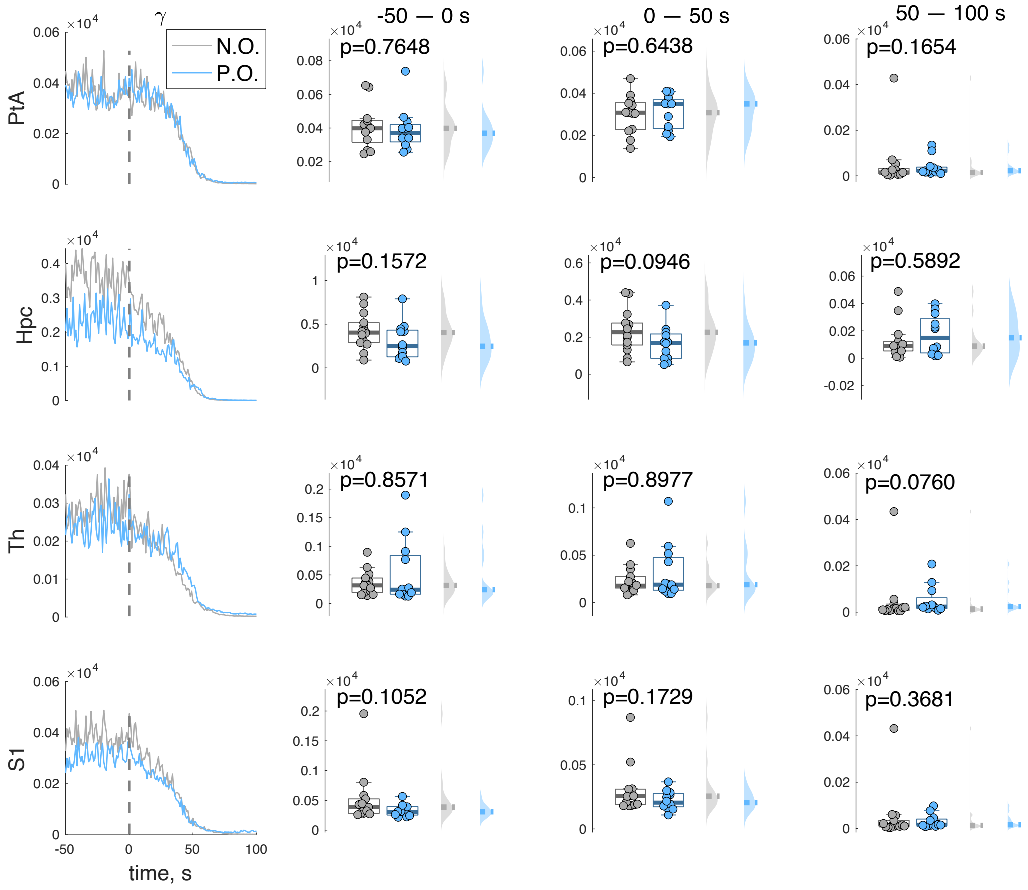


Figure S4. *Changes in the gamma band LFP activities following anoxia onset. The power of gamma (γ) activities in LFP records from the parietal association cortex (PtA), hippocampus (HpC), thalamus (Th), and primary somatosensory cortex (S1) was computed before and after cessation of brain oxygenation (t=0, thick dashed grey line). Changes in gamma activity were not significant between groups of rodents with positive (blue signals) or negative (grey signals) outcomes, regardless of the brain region or period examined. Thick lines in the box plots indicate median value, edges indicate the 25th to 75th percentiles, and whiskers extend up to 1.5 times the interquartile range from the edges.*

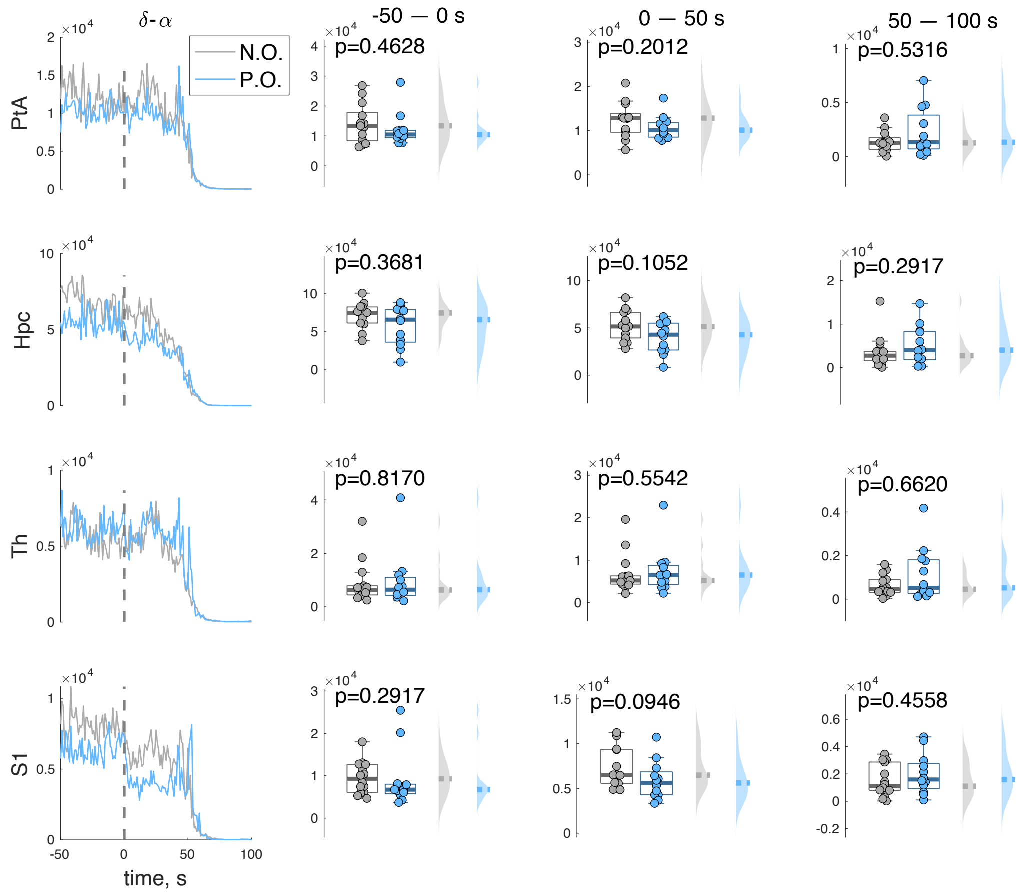


Figure S5. *Changes in the delta-alpha band LFP activities following anoxia onset. The power of delta-alpha (α–δ) activities in LFP records from the parietal association cortex (PtA), hippocampus (HpC), thalamus (Th), and primary somatosensory cortex (S1) was computed before and after cessation of brain oxygenation (t=0). Changes in alpha-delta activity were not significant between groups of rodents with positive (blue signals) or negative (grey signals) outcomes, regardless of the brain region or period examined. Thick lines in the box plots indicate median value, edges indicate the 25th to 75th percentiles, and whiskers extend up to 1.5 times the interquartile range from the edges.*

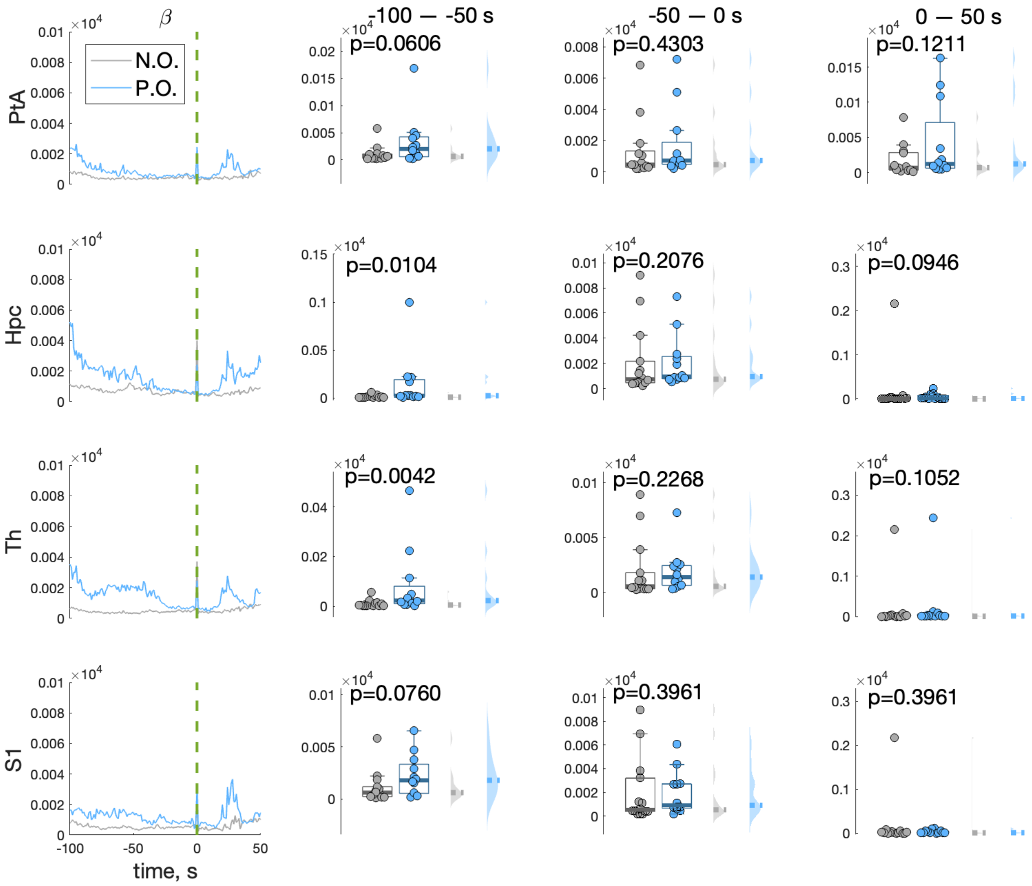


Figure S6. *Beta activities at the time of resuscitation. The power of beta (β) activities in LFP records from the parietal association cortex (PtA), hippocampus (HpC), thalamus (Th), and primary somatosensory cortex (S1) was computed before and after restoration of brain oxygenation (t=0, thick dashed green line). The power of beta activities, in each brain region, was higher (reaching significance only in HpC and Th) in the -100 – -50 s period preceding the start of the recue protocol in the rodents whose outcome was positive (blue signals). A rebound of activity during the 50 s following the resumption of ventilation was also observed in the P.O. group, although this increase remained not significant. Thick lines in the box plots indicate median value, edges indicate the 25th to 75th percentiles, and whiskers extend up to 1.5 times the interquartile range from the edges.*

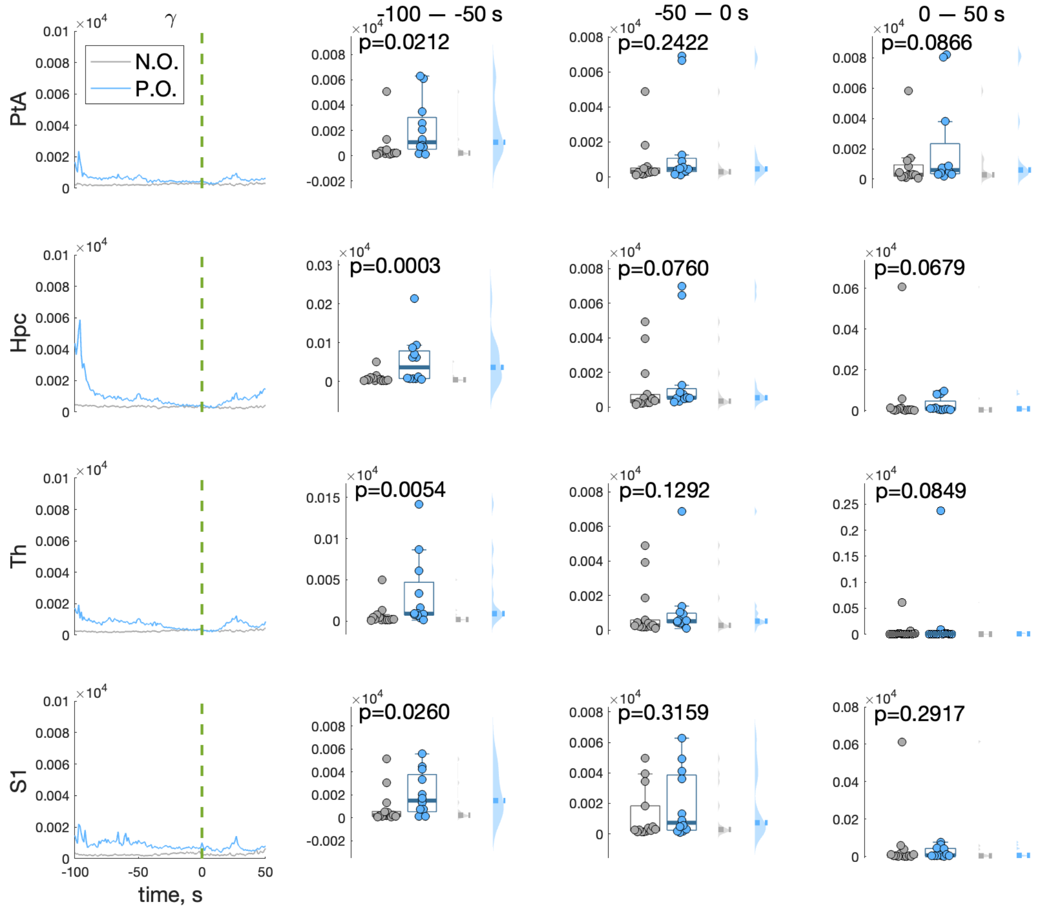


Figure S7. *Gamma activities at the time of resuscitation. The power of gamma (γ) activities in LFP records from the parietal association cortex (PtA), hippocampus (HpC), thalamus (Th), and primary somatosensory cortex (S1) was computed before and after restoration of brain oxygenation (t=0, thick dashed green line). The power of gamma activities in the -100 – -50 s period preceding restoration of brain oxygenation was higher in each recorded brain region in rodents whose outcome was positive (blue signals) as compared to rodents with negative outcomes (grey lines). A rebound of activity in the P.O. group during the 50 s following resuscitation was also observed in surviving rodents, although this increase remained not significant. Thick lines in the box plots indicate median value, edges indicate the 25th to 75th percentiles, and whiskers extend up to 1.5 times the interquartile range from the edges.*

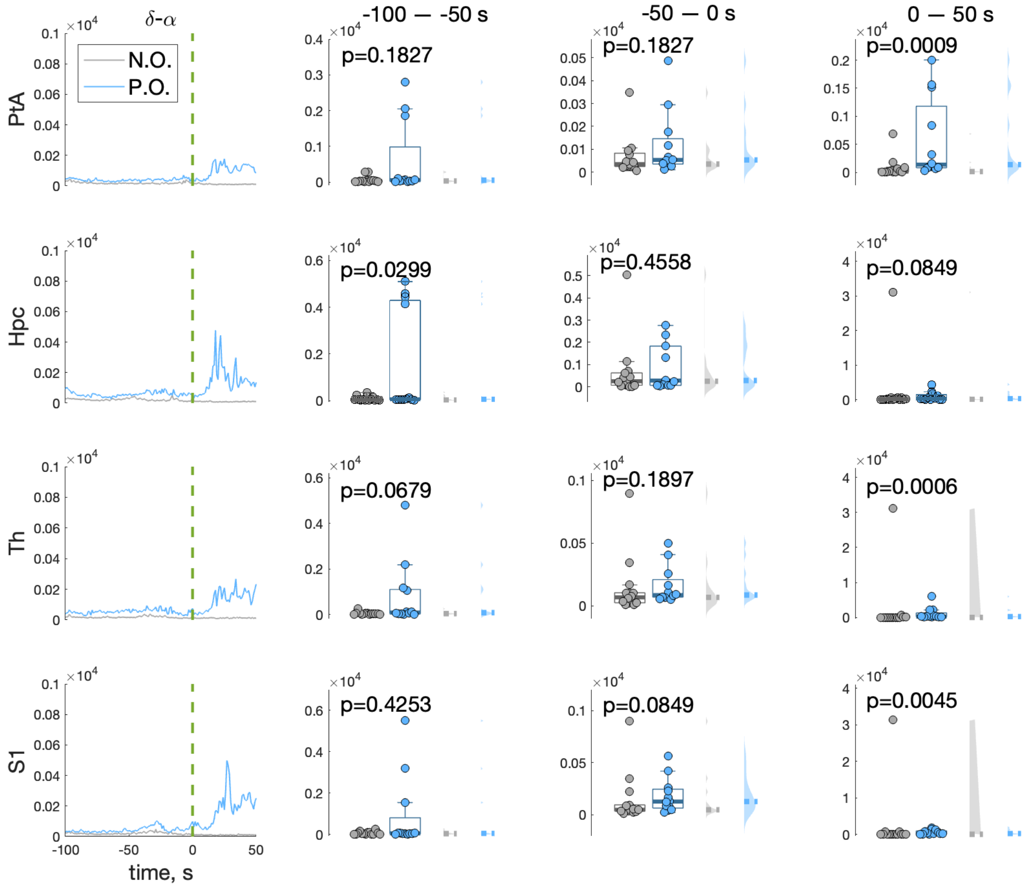


Figure S8. *Delta-alpha activities at the time of resuscitation. The power of delta-alpha (δ–α) activities in LFP records from the parietal association cortex (PtA), hippocampus (HpC), thalamus (Th), and primary somatosensory cortex (S1) was computed before and after restoration of brain oxygenation (t=0, thick dashed green line). In contrast to the other frequency bands, the power of delta-alpha activities in the -100 – -50 s period preceding restoration of brain oxygenation was similar in both groups of rodents. However, the rebound of activities in the P.O. group following oxygen resumption was significant in all brain regions, except in HpC. Thick lines in the box plots indicate median value, edges indicate the 25th to 75th percentiles, and whiskers extend up to 1.5 times the interquartile range from the edges.*

*Table S1. Statistical tests performed on the brain-heart coupling estimations, during resuscitation. The couplings were computed as the Spearman correlation coefficients between the envelope of LFP power and cardiac sympathetic of parasympathetic activity (CSI, CPI), within 0-50s with respect to the restoring of mechanical ventilation (vent. on). Wilcoxon tests were performed comparing the couplings in positive vs negative outcome individuals*

| | δ–α LFP – Cardiac fast HRV (CPI) | β LFP – Cardiac fast HRV (CPI) | γ LFP – Cardiac fast HRV (CPI) |
|---|---|---|---|
| Brain region | Wilcoxon test | Wilcoxon test | Wilcoxon test |
| PtA | p=0.9783<br>Z=-0.0271 | p=0.8066<br>Z=0.2447 | p=0.7236<br>Z=0.3535 |
| Hpc | p=0.1051<br>Z=1.620 | p=0.2687<br>Z=1.1058 | p=0.6996<br>Z=0.3857 |
| Th | p=0.6416<br>Z=0.4653 | p=1<br>Z=0 | p=0.5654<br>Z=-0.5748 |
| S1 | p=0.7236<br>Z=0.3535 | p=0.1085<br>Z=-0.5748 | p=0.2888<br>Z=1.0606 |
| | δ–α LFP – Cardiac slow HRV (CSI) | β LFP – Cardiac slow HRV (CSI) | γ LFP – Cardiac slow HRV (CSI) |
| | Wilcoxon test | Wilcoxon test | Wilcoxon test |
| PtA | p=0.8490<br>Z=-0.1903 | p=0.6438<br>Z=-0.4623 | p=0.5138<br>Z=0.6528 |
| Hpc | p=0.3413<br>Z=0.9515 | p=0.7772<br>Z=0.2828 | p=0.6251<br>Z=0.4886 |
| Th | p=0.3111<br>Z=1.0127 | p=0.5654<br>Z=-0.5748 | p=0.4598<br>Z=-0.7390 |
| S1 | p=0.6438<br>Z=0.4623 | p=0.5679<br>Z=0.5711 | p=0.2210<br>Z=1.2238 |

*Bold indicates significance, defined in Wilcoxon test as p<0.05 (Monte Carlo p) and in Chi-square test as p<0.0125 (Bonferroni corrected p). PtA: parietal association cortex, Hpc: hippocampus, Th: thalamus, S1: primary somatosensory cortex.*

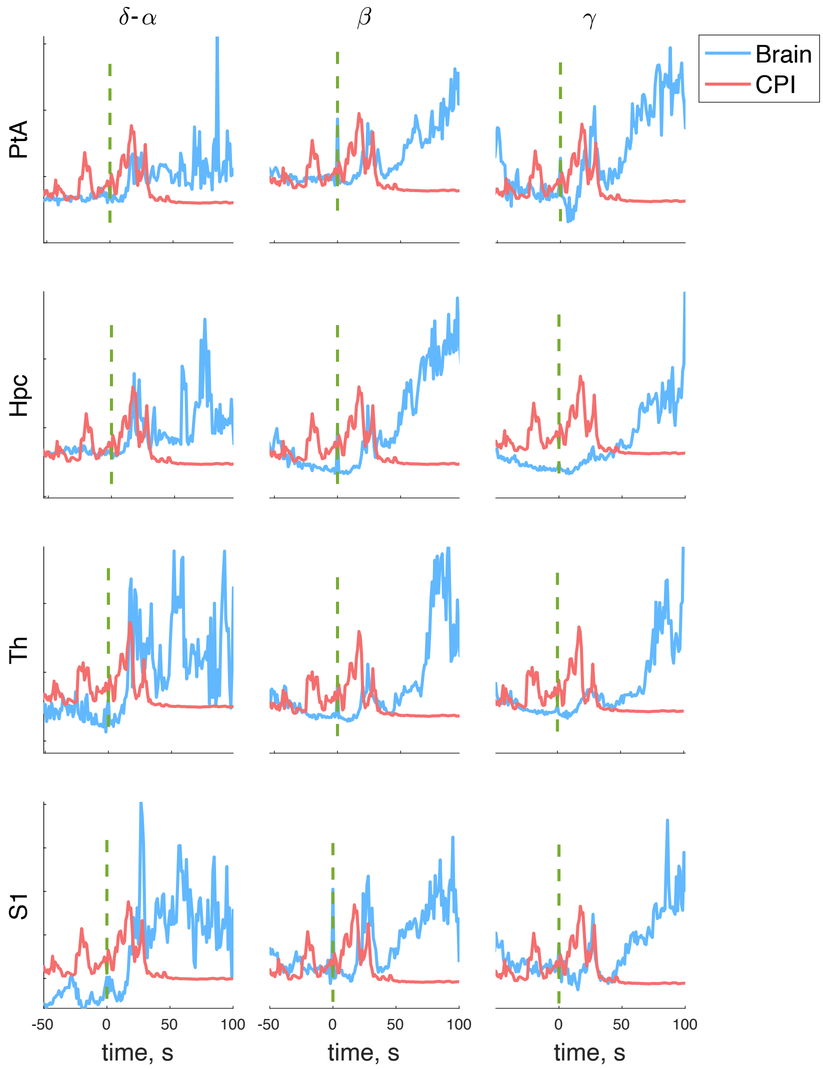


*Figure S9. Multi-site LFP and cardiac vagal responses to resuscitation. Group median, z-score normalized, changes in LFP power (blue lines), together with the temporal evolution of cardiac parasympathetic index (CPI, red lines), in the 100 s period after the return of brain oxygenation (t=0, thick dashed green line). Graphs correspond to the positive outcome group of rodents.*
